# Re-mining serum proteomics data reveals extensive post-translational modifications upon Zika and dengue infection

**DOI:** 10.1101/2022.06.02.494520

**Authors:** Kristina Allgoewer, Hyungwon Choi, Christine Vogel

## Abstract

Zika virus (ZIKV) and dengue virus (DENV) are two closely related flaviviruses with similar symptoms; understanding differences in their molecular impact on the host is therefore of high interest. Viruses interact with the host’s post-translational modifications, inducing changes visible in serum. As modifications are diverse and of low abundance, they typically require additional sample processing which is not feasible for large cohort studies. Therefore, we tested the potential of next-generation proteomics data in its ability to prioritize specific modifications for later targeted analysis. We re-mined published mass spectra from 122 unenriched serum samples from ZIKV and DENV patients for the presence of phosphorylated, methylated, oxidized, glycosylated/glycated, sulfated, and carboxylated peptides. We identified 272 modified peptides with significantly differential abundance in ZIKV and DENV patients. Amongst these, methionine-oxidized peptides from apolipoproteins and glycosylated peptides from immunoglobulin proteins were more abundant in ZIKV patient serum and generate hypotheses on the potential roles of the modification in the infection. The results demonstrate how data-independent acquisition techniques can help prioritize future analyses of peptide modifications.

## Introduction

Zika virus (ZIKV) and dengue virus (DENV) belong to the Flavivirus genus of the Flaviviridae family and are closely related. ZIKV and DENV cause a similar immune response in the host. While most infections are asymptomatic, symptoms can range from mild body pain to life-threatening fever (CDC 2020; Wen et al. 2017; Ngono and Shresta 2019). They overlap in their geographical distributions and can be transmitted by the same types of mosquitoes, *Aedes aegypti* and *Aedes albopictus* (CDC 2019; Ngono and Shresta 2019; CDC 2014). While the 2015 to 2017 outbreak in the Americas has waned, Zika cases have been surging in other places of the world such as India (Pierson and Diamond 2020; Sharma 2021; Bhargavi and Moa 2020). DENV infections are similarly widespread, resulting in ∼96 million symptomatic cases per year (World Health Organization 2020).

Viruses frequently interact with the cellular system of post-translational modifications (PTMs), changing host and viral proteins that may directly support the infection or modulate the antiviral response (Hu et al. 2020; Kumar et al. 2020). For example, DENV and ZIKV prevent STAT2 phosphorylation, which is required for the antiviral interferon response (Mazzon et al. 2009). Host proteins involved in the regulation of modifications can be potential drug targets that limit viral replication (Kumar et al. 2020). Common modifications in blood proteins are N- and O-glycosylation, phosphorylation, acylation/alkylation (e.g. acetylation, formylation, methylation, and pyroglutamylation), sulfation, hydroxylation, carboxylation, disulfide bridges, and proteolytic processing such as N- or C-terminal truncation (Schaller et al. 2008). Blood PTM signatures are emerging as biomarkers in clinical settings, e.g. for glycated hemoglobin, which is elevated in diabetes patients, and the phosphorylation status of vasodilator stimulated phosphoprotein to assess platelet reactivity (Mnatsakanyan et al. 2018).

However, modifications are challenging to analyze as they are highly diverse, with >400 known modifications (Aebersold et al. 2018), and typically of low abundance. Techniques for PTM-specific enrichment are cost and labor intensive (Doll and Burlingame 2015) which is often prohibitive for large sample numbers. Therefore, we tested the ability of existing data to help with prioritization of highly promising modifications for future analyses. Specifically, we examined a published mass spectrometry data set from 62 dengue and Zika patient serum samples (Allgoewer et al. 2021) (Figure 1A) which had been acquired in data-independent acquisition (DIA) mode for protein abundance analysis. DIA methods are well suited for such analysis as they generate fragmentation spectra for all ions captured in the full scan and contain all their ion chromatograms, providing the quantification abilities of targeted analyses with an expanded coverage of proteins and enabling the re-mining of existing data (Rosenberger et al. 2014; Faria et al. 2017). In addition, DIA has been shown to detect and quantify modifications reproducibly without prior enrichment (Goetze et al. 2020). Testing six PTM-specific libraries, we identified >1,800 modification events across eight different types of modifications of which methionine-oxidation, methylation, and glycosylation/glycation were most abundant and tested their differential abundance in the Zika and dengue patient serum samples.

**Figure 1:**
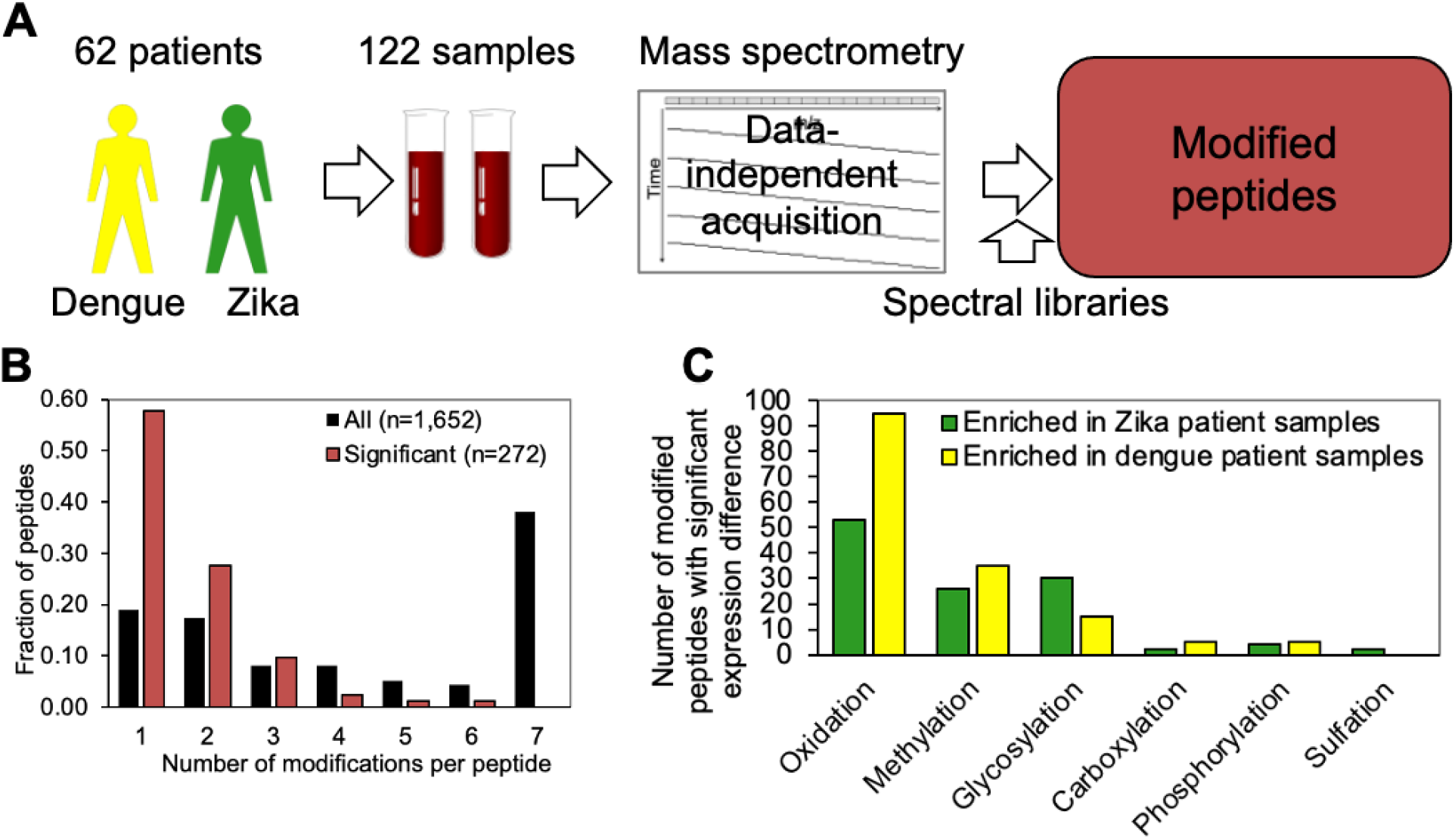
Re-mining serum proteomic data for modified peptides. **A)** We analyzed 122 serum samples from 62 patients from Trinidad, diagnosed with dengue and Zika as published in Allgoewer (2021). **B)** Most peptides observed and with significant expression differences had one modification, but some had more. **C)** Number of modified peptides with significant differential abundance between dengue and ZIKV patients. Modified peptides with significantly higher abundance (q-value ≤ 0.05, un-paired t-test) in dengue and ZIKV patients are shown in yellow and green, respectively. No acetylated and formylated peptides with significant differential abundance were identified.

## Methods and Materials

### Data and analysis

We obtained the RAW files from Allgoewer et al. (2021). The data had been collected in data-independent acquisition mode, comprising 124 patient and 20 quality control samples. RAW files were converted into the Spectronaut file format HTRMS (MS1/MS2 to centroid). All samples were analyzed using Spectronaut 14 against the project-specific spectral libraries with template fragments being used as the preferred fragment source for decoy generation. The remaining analysis settings remained in default. Two samples failed in their data acquisition and were removed from the analysis. False discovery rate (FDR), based on mProphet (Reiter et al. 2011), was calculated run-wise at the peptide precursor and experiment-wide at the protein group level and filtered for 1% on both levels. Modified peptides were extracted pooling different charge states.

### Spectral library generation

We prepared spectral libraries based on data acquired in data-dependent acquisition mode from pooled and fractionated serum samples of Zika and dengue patients (Allgoewer et al. 2021) using the Pulsar search engine within Spectronaut 14. Settings included Trypsin/P digest, peptide length of 7 to 52 amino acids and up to two missed cleavages. The FASTA file (human) was downloaded from UniProt (02/15/2018) and contained 93,798 entries including protein isoforms. Spectral libraries allowed for the following variable modifications: carboxylation, formylation, glycosylation (monohexose, dihexose, hexosamine, acetylhexosamine), methylation (monomethylation, dimethylation), phosphorylation, and sulfation (**Table S1**). In addition, acetylation (N-term) and oxidation (methionine) were set as variable modifications in each library. Carbamidomethylation of cysteine was set as fixed modification. Pulsar performs by default an internal mass calibration with optimized mass tolerance prediction for precursor and fragment ions. The FDR was calculated by Pulsar for peptide-spectrum matches, peptides, and protein groups and filtered for 1% at all three levels.

### Extraction of differentially abundant modified peptides

To identify differentially abundant modifications in serum of Zika and dengue patients, differential abundance was tested for modified peptides using the feature in Spectronaut 14 (two-sample t-test). We further processed the results to normalize for the abundance of the corresponding unmodified peptides. To do so, we subtracted the log base 2 fold change of the unmodified peptide from the log base 2 fold changes of modified peptide. We calculated q-values using the normalized log base 2 fold changes, the standard errors, and degrees of freedom (Benjamini and Hochberg 1995). Modified peptides without corresponding unmodified identifications were removed. Volcano plots were generated in R using the EnhancedVolcano() function.

### Protein function enrichment

Protein function information was retrieved from the PANTHER Classification System using the category “PANTHER protein class” (Mi et al. 2019; Mi et al. 2021). Actin-related proteins and non-motor actin binding proteins were summarized under the protein class parent “Cytoskeletal Protein”. Serine Proteases, Metalloproteases, and Proteases were summarized under the protein class parent “Protease”. Marimekko charts (mosaic plots) were generated in R using the ggplot() function. The size of each segment represents the number of unique modified peptides in each protein class and category. P-values for under- or overrepresentation were calculated using the UCLA Graeber Lab hypergeometric calculator (Graeber Lab 2009).

## Results

Using our workflow (**Figure 1A**), we generated spectral libraries specific to 12 different types of modifications (**Figure S1A**) and identified 1,652 modified peptides containing 1,806 different modifications (**Table 1, Table S5, Figure S1B**). The most common modification category was methionine-oxidation, followed by methylation, glycosylation/glycation, carboxylation, phosphorylation, sulfation, and acetylation(**Figure 1B, Figure S1C, Table S4**).

**Table 1.** Identified modifications. We identified 1,652 modified peptides containing 1,806 modifications. Modified residues are indicated using the one-letter amino acid code. For 1,122 modified peptides, the corresponding unmodified peptide could be identified, and fold changes were adjusted accordingly. We observed 272 modified peptides with significant differential abundance, with 117 and 155 being more abundant in Zika and dengue patients, respectively (**Table S6**).

|  |  |  | Modified peptides |  | Modified peptides significantly enriched in |  |
| --- | --- | --- | --- | --- | --- | --- |
| Modification | Modifications identified across all samples | Modified residues | Identified across all samples | With unmodified counterpart | ZIKV patients | dengue patients |
| **Oxidation** | 860 | M | 770 | 533 | 53 | 95 |
| **Methylation** | 447 | K, R | 426 | 296 | 26 | 35 |
| **Glycosylation/ Glycation** | 270 | K, N, R, S, T, W, Y | 257 | 175 | 30 | 15 |
| **Carboxylation** | 103 | D, E, K, W | 91 | 66 | 2 | 5 |
| **Phosphorylation** | 61 | STY | 53 | 30 | 4 | 5 |
| **Sulfation** | 33 | Y | 28 | 14 | 2 | 0 |
| **Acetylation (N-term)** | 29 | A, D, E, I, L, M, S, T, V | 24 | 5 | 0 | 0 |
| **Formylation (N-term)** | 3 | E, R, V | 3 | 3 | 0 | 0 |
| **Total** | **1806** |  | **1652** | **1122** | **117** | **155** |

The distribution of modifications amongst amino acid residues (**Figure S1C, Table S4**) was consistent with the literature, except for glycosylation: 37% of modifications were located on lysine and arginine, which are most common for glycation, which is often non-enzymatic (Baldensperger et al. 2020). We observed a surprising dominance of O-linked glycosylation events on serine, threonine, and tyrosine (47%), as serum proteins are reportedly predominantly N-linked glycosylated (Sun et al. 2018). C-linked glycosylation was generally rare, and we observed 11 peptides with this modification. Five of these contained the Trp-X-X-Trp consensus sequence (Krieg et al. 1998; Chauhan et al. 2013) and comprised several components of the Complement system which have been previously described as C-mannosylated (Hofsteenge et al. 1999) (**Table S3**).

**Figure 1C** shows the 272 modified peptides which differed significantly in their abundance between DENV and ZIKV patients (q-value ≤0.05)(**Table S6**). We observed more peptide oxidation, methylation, phosphorylation, and carboxylation in DENV than in ZIKV patients and more glycosylation/glycation and sulfation in ZIKV patients than in DENV patients. We identified no significant differences for acetylation and formylation.

### Methionine oxidation

We identified 148 methionine-oxidized peptides with significantly different abundance (q ≤ 0.05, **Table 1, Figure S2A**). While methionine-oxidation can result from sample preparation (Potgieter et al. 1997; Liu et al. 2013), all samples were processed similarly and evidence from literature supports the validity of the observed modification events.

For example, methionine oxidized peptides that were more abundant in ZIKV patients were significantly enriched for apolipoproteins (p = 0.0002, hypergeometric test, **Figure 2A/B**). The oxidation sites in the peptides from APOA1 (residue 136) and APOA2 (residue 49) have been described elsewhere (Pankhurst et al. 2003). In APOA1, methionine oxidation may affect structure and stability as well as impair the transport of cholesterol by inducing amyloid fibril formation. In the formation of infectious viral particles, host apolipoproteins play a role similar to Flaviviridae secretory proteins Erns and NS1 (Fukuhara et al. 2017), while APOE may be a target of the DENV capsid protein (Faustino et al. 2014).

**Figure 2:**
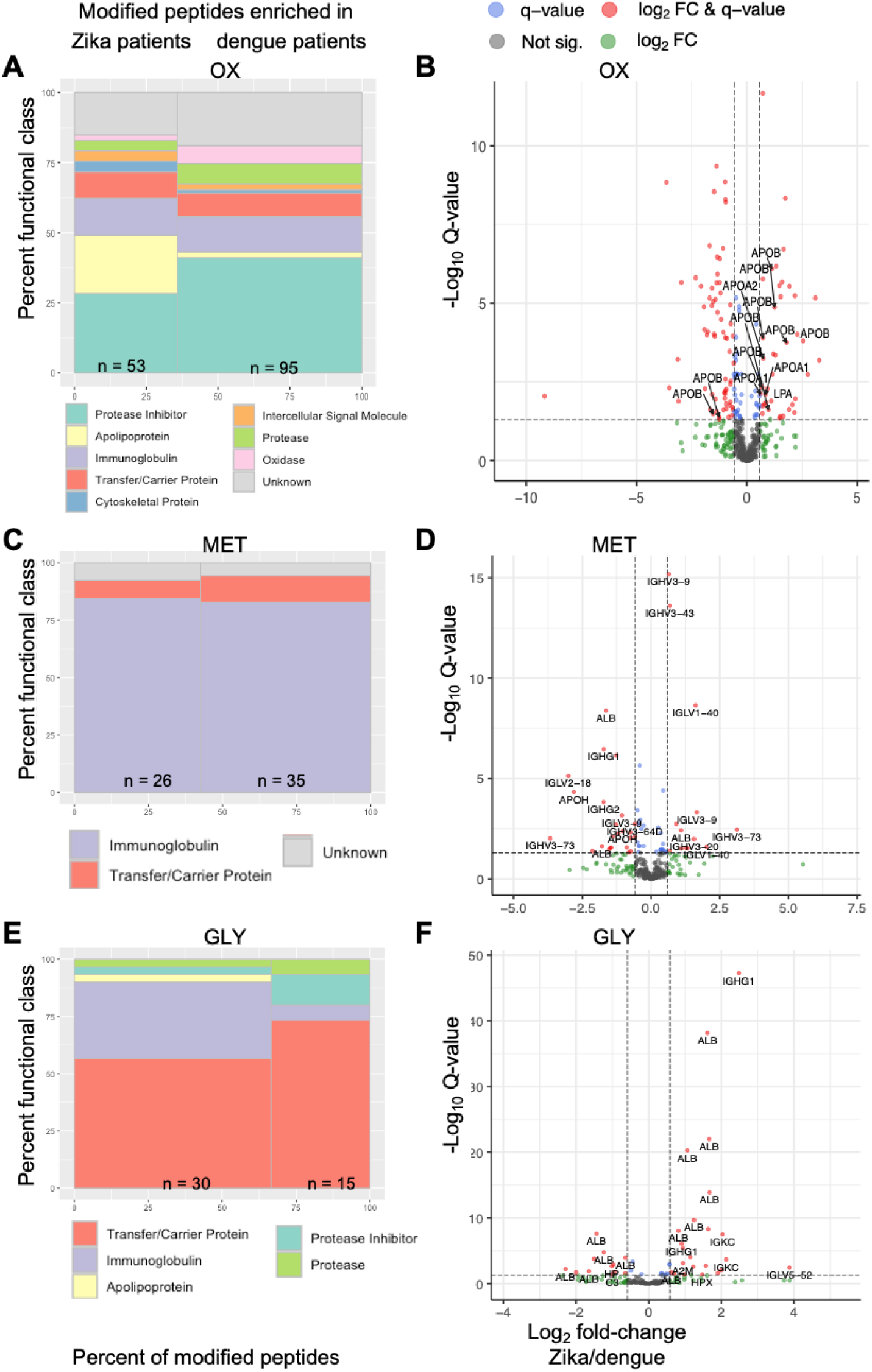
Function enrichment and abundance of the three most common modification types. Panels show results for oxidation (OX), methylation (MET), and glycosylation/glycation (GLY) in **A/B, C/D**, and **E/F**, respectively. Panels **A, C**, and **E** show the function enrichment obtained from the PANTHER classification system. Apolipoproteins and immunoglobulins are significantly overrepresented amongst methylated peptides (**A**) and glycosylated peptides (**E**), respectively (p-value ≤ 0.05) Panels **B, D**, and **F** show the volcano plots of fold-change (Zika/dengue) and q-value as a significance measure. q-value ≤ 0.05 (blue), absolute fold change (FC) ≥1.5 (green), beyond both thresholds (red), below both thresholds (grey). Panel **B** highlights apolipoproteins which were significantly overrepresented amongst methionine-oxidized peptides that are more abundant in serum from ZIKV patients.

Further, six methionine oxidized fibrinogen peptides were more abundant in ZIKV patients. Methionine oxidation in fibrinogen can change protein structure and impair aggregation and coagulation (Weigandt et al. 2012; White et al. 2016; Pederson and Interlandi 2019). The known modification sites were consistent with one of the modifications we identified in Fibrinogen A (residue 495) which was significantly less abundant in ZIKV patients.

All eleven methionine oxidized Alpha-2-Macroglobulin (A2M) peptides with significant differential abundance were more abundant in DENV patients (**Figure S2C**). The oxidation of A2M is known to alter the binding ability of cytokines, with possible implications for repair mechanisms during inflammation (Wu et al. 1998). Five methionine oxidized Alpha-1-Antitrypsin (SERPINA1) peptides were less abundant in ZIKV patients (**Figure S2D**). Methionine oxidation in SERPINA1 can influence protein structure (Griffiths and Cooney 2002) and regulate protein activity at inflammation sites (Taggart et al. 2000; Li et al. 2009).

### Methylation

We identified 61 methylated peptides with significantly different abundance in DENV and ZIKV patients (q-value ≤ 0.05, **Table 1, Figure 2C/D**). The methylated peptides were enriched in transfer/carrier proteins and immunoglobulins (p-value≤ 0.05). All three methylated peptides belonging to immunoglobulin class G were significantly more abundant in DENV patients. In addition, we found extensive methylation of Serum Albumin. While most protein methylation events have been described for histone proteins (Liu et al. 2013), Arginine monomethylation was found to be one of the most abundant modifications in the serum of cancer patients (Gu et al. 2016). Gu et al. (2016) speculate that Arginine methylation could either be catalyzed in the bloodstream or be released from intracellular compartments, since various Arginine methyltransferases can occur in plasma (Liu et al. 2007; Sennels et al. 2007). Key platelet proteins are known to be Arginine methylated, which may promote aggregation in response to thrombin and collagen (Marsden et al. 2021). In mouse brain tissue, Arginine methylation has been detected in transporters and vesicle proteins as well as in ion channels and receptors, all of which are involved in synaptic transmission (Guo et al. 2014).

### Glycosylation and glycation

The third most frequently identified modification was glycation, where two thirds of the modified peptides were more abundant in ZIKV patients (q-value ≤ 0.05, **Table 1, Figure 2E/F**). The 46 modifications included 41 monohexoses, 4 hexosamines, and one acetylhexosamine. Immunoglobulins were significantly overrepresented among peptides that are more abundant in ZIKV patients (p≤ 0.05). We observed 23 modifications on Lys/Arg residues, rendering them more likely to be glycations (Baldensperger et al. 2020). Three of the remaining 23 modification sites were N-linked glycosylations; 20 were O-linked of which many were more abundant in ZIKV patients (**Figure S3A**).

A large number of glycosylated/glycated peptides derived from serum albumin (27) and immunoglobulin components (11)(**Figures S3B/C**). Glycation of serum albumin influences its conformation and function and leads to a shorter half-life, with the primary glycation site being Lys-525 (Jones et al. 1983; Shaklai et al. 1984). We observed this modification in two of the identified peptides. Glycation of immunoglobulins can occur *in vitro* and *in vivo*: while no impact on function could be detected in human antibodies (Goetze et al. 2012), activity loss through glycation has been reported in some studies on therapeutic antibodies (Wei et al. 2017). All three glycosylation/glycation events that were detected on IgG peptides were more abundant in ZIKV patients. Alterations in immunoglobulin glycosylations are also associated with aging, disease, and inflammation (Gudelj et al. 2018; Karsten et al. 2012). Afucosylation of maternal anti-DENV IgGs has been shown to influence susceptibility of infants to symptomatic disease (Thulin et al. 2020).

Finally, we observed a modified haptoglobin peptide, i.e. a glycation event (Lys, monohexose) that was significantly less abundant in ZIKV patients. This specific modification (glycated Lys-141) has been described as a potential biomarker for the early diagnosis of type 2 diabetes (Spiller et al. 2017).

### Other modifications

We identified phosphorylated peptides that differed significantly in their abundance between dengue and ZIKV patients (q-value ≤0.05). Phosphoproteins and phosphorylated sites are known to be upregulated in DENV infected cells compared to controls (Miao et al. 2019). The phosphorylated sites included residues in serum albumin, Fibronectin, S100-A9, and in Kininogen. Phosphorylated Fibronectin has been shown to enhance cell attachment (Yalak et al. 2019), and fibronectin was reported to be elevated in the serum of pregnant ZIKV patients (Foo et al. 2017). Altered expression of the inflammatory protein S100-A9 is associated with recurrent pregnancy loss (Nair et al. 2013). Both its intracellular and extracellular function was shown to be regulated by its phosphorylation in the specific site observed in our data, impacting the proinflammatory function of neutrophils (Schenten et al. 2018). In its high molecular weight form, Kininogen also has proinflammatory functions, and was shown to be downregulated in dengue patients (Oehmcke-Hecht and Köhler 2018; Ray et al. 2012). We also identified phosphorylation in Fibrinogen Alpha. Phosphorylation of fibrinogen has been suggested to alter its function, increase fiber diameters, and potentially prevent bleeding (Martin et al. 1992; de Vries et al. 2020).

We further identified 7 carboxylated peptides which differed significantly in their abundance between DENV and ZIKV patients (q-value ≤ 0.05, **Table 1, Figure S4C**). Carboxylation most commonly occurs on Glu residues and is associated with blood clotting, extraosseous calcification, and bone growth (Lee et al. 2011). Lys carboxylation has been shown to modulate the activity of enzymes (Li et al. 2010; Jimenez-Morales et al. 2013). In human blood cells, increased protein carboxylation has been observed after treatment with catechol (Bukowska et al. 2015).

We observed two sulfation events in different immunoglobulin regions (IGKC and IGHV4-34) which are more abundant in ZIKV patients. Both sulfation modifications are located on Tyr residues, which has been shown to alter immunoglobulin potency and contributes to the diversity of the humoral immune response (Choe et al. 2003).

Finally, we observed N-terminally acetylated and N-terminally formylated peptides, but without significant differences in abundance between ZIKV and DENV patients (**Table 1**). Acetylation is the second most frequent intracellular modification (Khoury et al. 2011). The low number of identifications in our data, may indicate the need for special enrichment methods, which are often deemed essential for the analysis of N-terminal modifications (Yeom et al. 2017; Chang et al. 2021). The acetylated peptides belong to Haptoglobin-Related Protein, immunoglobulin fragment IGHV3OR16-9, Peptidyl-Prolyl Cis-Trans Isomerase A, Hemoglobin Subunit Alpha 1, and Hemoglobin Subunit Beta. Altered protein acetylation may play a role in certain diseases such as rheumatoid arthritis (Arito et al. 2015).

## Discussion

Analysis of protein modifications typically requires selective enrichment of modified peptides (Larsen et al. 2006) - which can be labor and cost intensive if conducted at a large scale and is not possible for all types of modifications. Therefore, we tested the potential of re-mining DIA data to identify modifications in a cohort of serum samples without prior enrichment. We identified >1,800 modification events across data from Zika and dengue patients (**Table S5**), 272 of which were significantly different between the two types of infection (**Table S6**). Literature supports the validity of the results in their potential to generate new hypotheses.

Future work might investigate methionine oxidation: the 148 oxidized peptides with significantly differential abundances in serum from Zika and dengue patients suggest true biological relevance. As non-enzymatic reactions during sample preparation can affect the results (Potgieter et al. 1997; Morand et al. 1993), future experiments would require oxidation of all methionine with isotopically labeled hydrogen peroxide to distinguish between true and false positive oxidation events (Liu et al. 2013).

Specifically, we observed an enrichment in methionine-oxidation on apolipoproteins with higher abundance in Zika samples. Host apolipoproteins play a role in Flaviviridae infection (Fukuhara et al. 2017); for example, Apolipoprotein E (APOE) interacts with the DENV capsid protein (Faustino et al. 2014). Some APOE genotypes are associated with microcephaly and neurocognitive disorders (Oriá et al. 2005; Geffin et al. 2017; Braga et al. 2010; Kuroda et al. 2007). Further, methionine oxidation of Apolipoprotein A-I can induce amyloid fibril formation, impairing transport of cholesterol (Wong et al. 2010; Shao et al. 2008). Host cholesterol levels, in turn, support virus replicative complex formation, virus assembly and egress (Li et al. 2013)(Li et al. 2013; Osuna-Ramos et al. 2018). Finally, hepatitis C virus oxidizes APOB, resulting in degradation by the 20S proteasome (Wang et al. 2021).

While phosphorylation is typically the most abundant PTM in eukaryotic cells (Khoury et al. 2011), it is less abundant in the extracellular space (Chen et al. 2017). Highly active phosphatases in blood further challenge identification. The fact that we still detected 9 significantly different phosphorylation events suggests that phosphorylation is another modification worth exploring in future experiments. However, enrichment and the use of phosphatase inhibitors during sample preparation will be essential (Chen et al. 2017).

Another modification with highly promising results in our data was glycosylation/glycation, for which we observed 45 modified peptides, in particular from immunoglobulin proteins, with significant abundance differences between Zika and dengue patient serum. Glycosylations in immunoglobulin components are associated with aging, disease, inflammation, pathophysiology and immune suppression (Gudelj et al. 2018; Karsten et al. 2012; Bovenkamp et al. 2016). Changes in glycosylation patterns of IgG have been observed in patients with autoimmune disease such as rheumatoid arthritis (Parekh et al. 1985; Maverakis et al. 2015). Consistent with our observations, specific subclasses of neutralizing antibodies from ZIKV patients require glycosylation for binding to ZIKV envelope proteins (Henderson et al. 2020) - generating hypotheses on the role of this modification in this infection.

Further, we observed substantial glycation, in particular of serum albumin. Glycation of serum albumin, however, is known to cause a shorter half-life (Jones et al. 1983; Shaklai et al. 1984) - and our data contained the primary glycation site responsible for this effect. Serum albumin glycation is a biomarker for diabetes mellitus and a potential predictor for renal, cerebro- and cardio-metabolic diseases (Giglio et al. 2020). Our results therefore motivate the future analysis of the structure and function of glycoforms, e.g. by hydrophilic interaction liquid chromatography with fluorescence detection (de Haan et al. 2020).

## Acknowledgements

CV acknowledges funding by the US National Institutes of Health (R35GM127089 and 75N93019C00052/NH/NIH HHS/United States). KA acknowledges funding by the predoctoral fellowship by the German Academic Exchange Service (DAAD, “Jahresstipendium für Doktorandinnen und Doktoranden bei bi-national betreuten Promotionen”).

## References

Allgoewer, K., Maity, S., Zhao, A., Lashua, L., Ramgopal, M., Balkaran, B.N., Liu, L., Purushwani, S., Arévalo, M.T., Ross, T.M., Choi, H., Ghedin, E. & Vogel, C. (2021) New Proteomic Signatures to Distinguish Between Zika and Dengue Infections. Molecular & cellular proteomics: MCP 20: 100052.

Arito, M., Nagai, K., Ooka, S., Sato, T., Takakuwa, Y., Kurokawa, M.S., Sase, T., Okamoto, K., Suematsu, N. & Kato, T. (2015) Altered acetylation of proteins in patients with rheumatoid arthritis revealed by acetyl-proteomics. Clinical and Experimental Rheumatology 33(6): 877–886.

Aubry, F., Jacobs, S., Darmuzey, M., Lequime, S., Delang, L., Fontaine, A., Jupatanakul, N., Miot, E.F., Dabo, S., Manet, C., Montagutelli, X., Baidaliuk, A., Gámbaro, F., Simon-Lorière, E., Gilsoul, M., Romero-Vivas, C.M., Cao-Lormeau, V.-M., Jarman, R.G., Diagne, C.T., Faye, Oumar, Faye, Ousmane, Sall, A.A., Neyts, J., Nguyen, L., Kaptein, S.J.F. & Lambrechts, L. (2021) Recent African strains of Zika virus display higher transmissibility and fetal pathogenicity than Asian strains. Nature Communications 12(1): 916.

Baldensperger, T., Eggen, M., Kappen, J., Winterhalter, P.R., Pfirrmann, T. & Glomb, M.A. (2020) Comprehensive analysis of posttranslational protein modifications in aging of subcellular compartments. Scientific Reports 10(1): 7596.

Bantscheff, M., Lemeer, S., Savitski, M.M. & Kuster, B. (2012) Quantitative mass spectrometry in proteomics: critical review update from 2007 to the present. Analytical and Bioanalytical Chemistry 404(4): 939–965.

Bantscheff, M., Schirle, M., Sweetman, G., Rick, J. & Kuster, B. (2007) Quantitative mass spectrometry in proteomics: a critical review. Analytical and Bioanalytical Chemistry 389(4): 1017–1031.

Barber, K.W. & Rinehart, J. (2018) The ABCs of modifications. Nature chemical biology 14(3): 188–192.

Benjamini, Y. & Hochberg, Y. (1995) Controlling the False Discovery Rate: a Practical and Powerful Approach to Multiple Testing. Available from: /paper/Controlling-the-False-Discovery-Rate%3A-a-Practical-Benjamini-Hochberg/fcef2258a963f3d3984a486185ddc4349c43aa35 [Accessed 11 June 2020].

Bhargavi, B.S. & Moa, A. (2020) Global outbreaks of zika infection by epidemic observatory (EpiWATCH), 2016-2019. Global Biosecurity 2(1). Available from: http://jglobalbiosecurity.com/articles/10.31646/gbio.83/ [Accessed 18 September 2021].

Bovenkamp, F.S. van de, Hafkenscheid, L., Rispens, T. & Rombouts, Y. (2016) The Emerging Importance of IgG Fab Glycosylation in Immunity. The Journal of Immunology 196(4): 1435–1441.

Braga, L.W., Borigato, E.V.M., Speck-Martins, C.E., Imamura, E.U., Gorges, A.M.P., Izumi, A.P., Dantas, R.C. & Nunes, L.G.N. (2010) Apolipoprotein E genotype and cerebral palsy. Developmental Medicine and Child Neurology 52(7): 666–671.

Bukowska, B., Michałowicz, J. & Marczak, A. (2015) The effect of catechol on human peripheral blood mononuclear cells (in vitro study). Environmental Toxicology and Pharmacology 39(1): 187–193.

Burk, R.F., Hill, K.E., Motley, A.K., Winfrey, V.P., Kurokawa, S., Mitchell, S.L. & Zhang, W. (2014) Selenoprotein P and apolipoprotein E receptor-2 interact at the blood-brain barrier and also within the brain to maintain an essential selenium pool that protects against neurodegeneration. FASEB journal: official publication of the Federation of American Societies for Experimental Biology 28(8): 3579–3588.

CDC (2020) Dengue Virus. Available from: https://www.cdc.gov/dengue/index.html [Accessed 17 May 2021].

CDC (2019) Flaviviridae | Viral Hemorrhagic Fevers (VHFs). Available from: https://www.cdc.gov/vhf/virus-families/flaviviridae.html [Accessed 8 June 2021].

CDC (2014) Zika Virus. Available from: https://www.cdc.gov/zika/index.html [Accessed 17 May 2021].

Chang, C.-H., Chang, H.-Y., Rappsilber, J. & Ishihama, Y. (2021) Isolation of Acetylated and Unmodified Protein N-Terminal Peptides by Strong Cation Exchange Chromatographic Separation of TrypN-Digested Peptides. Molecular & Cellular Proteomics 20: 100003.

Chauhan, J.S., Rao, A. & Raghava, G.P.S. (2013) In silico Platform for Prediction of N-, O-and C-Glycosites in Eukaryotic Protein Sequences. PLOS ONE 8(6): e67008.

Chen, I.-H., Xue, L., Hsu, C.-C., Paez, J.S.P., Pan, L., Andaluz, H., Wendt, M.K., Iliuk, A.B., Zhu, J.-K. & Tao, W.A. (2017) Phosphoproteins in extracellular vesicles as candidate markers for breast cancer. Proceedings of the National Academy of Sciences 114(12): 3175–3180.

Cheng, L.C., Li, Z., Graeber, T.G., Graham, N.A. & Drake, J.M. (2018) Phosphopeptide Enrichment Coupled with Label-free Quantitative Mass Spectrometry to Investigate the Phosphoproteome in Prostate Cancer. Journal of Visualized Experiments : JoVE (138): 57996.

Choe, H., Li, W., Wright, P.L., Vasilieva, N., Venturi, M., Huang, C.-C., Grundner, C., Dorfman, T., Zwick, M.B., Wang, L., Rosenberg, E.S., Kwong, P.D., Burton, D.R., Robinson, J.E., Sodroski, J.G. & Farzan, M. (2003) Tyrosine sulfation of human antibodies contributes to recognition of the CCR5 binding region of HIV-1 gp120. Cell 114(2): 161–170.

Cox, J., Hein, M.Y., Luber, C.A., Paron, I., Nagaraj, N. & Mann, M. (2014) Accurate proteome-wide label-free quantification by delayed normalization and maximal peptide ratio extraction, termed MaxLFQ. Molecular & cellular proteomics: MCP 13(9): 2513–2526.

Doll, S. & Burlingame, A.L. (2015) Mass spectrometry-based detection and assignment of protein posttranslational modifications. ACS chemical biology 10(1): 63–71.

Eggo, R.M. & Kucharski, A.J. (2018) Expected Duration of Adverse Pregnancy Outcomes after Zika Epidemic. Emerging Infectious Diseases 24(1): 127–130.

Faria, S., Morris, C., Silva, A., Fonseca, M., Forget, P., S. Castro M. & Fontes, W. (2017) A Timely Shift from Shotgun to Targeted Proteomics and How It Can Be Groundbreaking for Cancer Research. Frontiers in Oncology 7.

Faustino, A.F., Carvalho, F.A., Martins, I.C., Castanho, M.A.R.B., Mohana-Borges, R., Almeida, F.C.L., Da Poian, A.T. & Santos, N.C. (2014) Dengue virus capsid protein interacts specifically with very low-density lipoproteins. Nanomedicine: Nanotechnology, Biology and Medicine 10(1): 247–255.

Foo, S.-S., Chen, W., Chan, Y., Bowman, J.W., Chang, L.-C., Choi, Y., Yoo, J.S., Ge, J., Cheng, G., Bonnin, A., Nielsen-Saines, K., Brasil, P. & Jung, J.U. (2017) Asian Zika virus strains target CD14+ blood monocytes and induce M2-skewed immunosuppression during pregnancy. Nature Microbiology 2(11): 1558–1570.

Fukuhara, T., Tamura, T., Ono, C., Shiokawa, M., Mori, H., Uemura, K., Yamamoto, S., Kurihara, T., Okamoto, T., Suzuki, R., Yoshii, K., Kurosu, T., Igarashi, M., Aoki, H., Sakoda, Y. & Matsuura, Y. (2017) Host-derived apolipoproteins play comparable roles with viral secretory proteins Erns and NS1 in the infectious particle formation of Flaviviridae. PLoS pathogens 13(6): e1006475.

Gao, J. & Xu, D. (2012) Correlation between posttranslational modification and intrinsic disorder in protein. Pacific Symposium on Biocomputing. Pacific Symposium on Biocomputing: 94–103.

Geffin, R., Martinez, R., de Las Pozas, A., Issac, B. & McCarthy, M. (2017) Apolipoprotein E4 Suppresses Neuronal-Specific Gene Expression in Maturing Neuronal Progenitor Cells Exposed to HIV. Journal of Neuroimmune Pharmacology: The Official Journal of the Society on NeuroImmune Pharmacology 12(3): 462–483.

Giglio, R.V., Lo Sasso, B., Agnello, L., Bivona, G., Maniscalco, R., Ligi, D., Mannello, F. & Ciaccio, M. (2020) Recent Updates and Advances in the Use of Glycated Albumin for the Diagnosis and Monitoring of Diabetes and Renal, Cerebro-and Cardio-Metabolic Diseases. Journal of Clinical Medicine 9(11): E3634.

Goetze, A.M., Liu, Y.D., Arroll, T., Chu, L. & Flynn, G.C. (2012) Rates and impact of human antibody glycation in vivo. Glycobiology 22(2): 221–234.

Goetze, S., Frey, K., Rohrer, L., Radosavljevic, S., Krützfeldt, J., Landmesser, U., Bueter, M., Pedrioli, P.G.A., Eckardstein, A. von & Wollscheid, B. (2020) Sensitive and reproducible determination of clinical HDL proteotypes. bioRxiv [Preprint]: 2020.07.09.191312.

Graeber Lab (2009) Hypergeometric p-value calculator. Available from: https://systems.crump.ucla.edu/hypergeometric/index.php [Accessed 26 May 2021].

Griffiths, S.W. & Cooney, C.L. (2002) Relationship between protein structure and methionine oxidation in recombinant human alpha 1-antitrypsin. Biochemistry 41(20): 6245–6252.

Gu, H., Ren, J.M., Jia, X., Levy, T., Rikova, K., Yang, V., Lee, K.A., Stokes, M.P. & Silva, J.C. (2016) Quantitative Profiling of Post-translational Modifications by Immunoaffinity Enrichment and LC-MS/MS in Cancer Serum without Immunodepletion. Molecular & Cellular Proteomics 15(2): 692–702.

Gudelj, I., Lauc, G. & Pezer, M. (2018) Immunoglobulin G glycosylation in aging and diseases. Cellular Immunology 333: 65–79.

Guo, A., Gu, H., Zhou, J., Mulhern, D., Wang, Y., Lee, K.A., Yang, V., Aguiar, M., Kornhauser, J., Jia, X., Ren, J., Beausoleil, S.A., Silva, J.C., Vemulapalli, V., Bedford, M.T. & Comb, M.J. (2014) Immunoaffinity enrichment and mass spectrometry analysis of protein methylation. Molecular & cellular proteomics: MCP 13(1): 372–387.

de Haan, N., Falck, D. & Wuhrer, M. (2020) Monitoring of immunoglobulin N-and O-glycosylation in health and disease. Glycobiology 30(4): 226–240.

Halim, A., Brinkmalm, G., Rüetschi, U., Westman-Brinkmalm, A., Portelius, E., Zetterberg, H., Blennow, K., Larson, G. & Nilsson, J. (2011) Site-specific characterization of threonine, serine, and tyrosine glycosylations of amyloid precursor protein/amyloid β-peptides in human cerebrospinal fluid. Proceedings of the National Academy of Sciences of the United States of America 108(29): 11848–11853.

Hartel, N.G., Chew, B., Qin, J., Xu, J. & Graham, N.A. (2019) Deep Protein Methylation Profiling by Combined Chemical and Immunoaffinity Approaches Reveals Novel PRMT1 Targets. Molecular & Cellular Proteomics 18(11): 2149–2164.

Henderson, E.A., Tam, C.C., Cheng, L.W., Ngono, A.E., Nguyen, A.-V., Shresta, S., McGee, M., Padgett, H., Grill, L.K. & Martchenko Shilman, M. (2020) Investigation of the immunogenicity of Zika glycan loop. Virology Journal 17(1): 43.

Hofsteenge, J., Blommers, M., Hess, D., Furmanek, A. & Miroshnichenko, O. (1999) The four terminal components of the complement system are C-mannosylated on multiple tryptophan residues. The Journal of Biological Chemistry 274(46): 32786–32794.

Howard, C.R. ed. (2005) Flaviviruses. Perspectives in Medical Virology 11: 13–51.

Hu, J., Zhang, L. & Liu, X. (2020) Role of Post-translational Modifications in Influenza A Virus Life Cycle and Host Innate Immune Response. Frontiers in Microbiology 11: 2156.

Jimenez-Morales, D., Adamian, L., Shi, D. & Liang, J. (2013) Lysine carboxylation: unveiling a spontaneous post-translational modification. Acta Crystallographica Section D: Biological Crystallography 70(Pt 1): 48–57.

Jones, I.R., Owens, D.R., Williams, S., Ryder, R.E., Birtwell, A.J., Jones, M.K., Gicheru, K. & Hayes, T.M. (1983) Glycosylated serum albumin: an intermediate index of diabetic control. Diabetes Care 6(5): 501–503.

Karsten, C.M., Pandey, M.K., Figge, J., Kilchenstein, R., Taylor, P.R., Rosas, M., McDonald, J.U., Orr, S.J., Berger, M., Petzold, D., Blanchard, V., Winkler, A., Hess, C., Reid, D.M., Majoul, I.V., Strait, R.T., Harris, N.L., Köhl, G., Wex, E., Ludwig, R., Zillikens, D., Nimmerjahn, F., Finkelman, F.D., Brown, G.D., Ehlers, M. & Köhl, J. (2012) Anti-inflammatory activity of IgG1 mediated by Fc galactosylation and association of FcγRIIB and dectin-1. Nature Medicine 18(9): 1401–1406.

Khoury, G.A., Baliban, R.C. & Floudas, C.A. (2011) Proteome-wide post-translational modification statistics: frequency analysis and curation of the swiss-prot database. Scientific Reports 1. Available from: https://www.ncbi.nlm.nih.gov/pmc/articles/PMC3201773/ [Accessed 15 March 2021].

Kim, J.-M., Seok, O.-H., Ju, S., Heo, J.-E., Yeom, J., Kim, D.-S., Yoo, J.-Y., Varshavsky, A., Lee, C. & Hwang, C.-S. (2018) Formyl-methionine as an N-degron of a eukaryotic N-end rule pathway. Science (New York, N.Y.) 362(6418).

Krasny, L. & Huang, P.H. (2021) Data-independent acquisition mass spectrometry (DIA-MS) for proteomic applications in oncology. Molecular Omics 17(1): 29–42.

Krieg, J., Hartmann, S., Vicentini, A., Gläsner, W., Hess, D. & Hofsteenge, J. (1998) Recognition Signal for C-Mannosylation of Trp-7 in RNase 2 Consists of Sequence Trp-x-x-Trp. Molecular Biology of the Cell 9(2): 301–309.

Kumar, R., Mehta, D., Mishra, N., Nayak, D. & Sunil, S. (2020) Role of Host-Mediated Post-Translational Modifications (PTMs) in RNA Virus Pathogenesis. International Journal of Molecular Sciences 22: 323.

Kuroda, M.M., Weck, M.E., Sarwark, J.F., Hamidullah, A. & Wainwright, M.S. (2007) Association of Apolipoprotein E Genotype and Cerebral Palsy in Children. Pediatrics 119(2): 306–313.

Larsen, M.R., Trelle, M.B., Thingholm, T.E. & Jensen, O.N. (2006) Analysis of posttranslational modifications of proteins by tandem mass spectrometry. BioTechniques 40(6): 790–798.

Lee, T.-Y., Lu, C.-T., Chen, S.-A., Bretaña, N.A., Cheng, T.-H., Su, M.-G. & Huang, K.-Y. (2011) Investigation and identification of protein γ-glutamyl carboxylation sites. BMC Bioinformatics 12(13): S10.

Li, Y., Kakinami, C., Li, Q., Yang, B. & Li, H. (2013) Human Apolipoprotein A-I Is Associated with Dengue Virus and Enhances Virus Infection through SR-BI. PLoS ONE 8(7). Available from: https://www.ncbi.nlm.nih.gov/pmc/articles/PMC3722190/ [Accessed 28 March 2021].

Li, Y., Yu, X., Ho, J., Fushman, D., Allewell, N.M., Tuchman, M. & Shi, D. (2010) Reversible Post-Translational Carboxylation Modulates The Enzymatic Activity Of N-Acetyl-L-Ornithine Transcarbamylase. Biochemistry 49(32): 6887–6895.

Li, Z., Alam, S., Wang, J., Sandstrom, C.S., Janciauskiene, S. & Mahadeva, R. (2009) Oxidized {alpha}1-antitrypsin stimulates the release of monocyte chemotactic protein-1 from lung epithelial cells: potential role in emphysema. American Journal of Physiology. Lung Cellular and Molecular Physiology 297(2): L388–400.

Liu, Huadong, Galka, M., Liu, X., Lin, Y., Pittock, P., Voss, C., Dhami, G., Li, X., Miyaji, M., Lajoie, G., Chen, B. & Li, S.S.-C. (2013) A method for systematic mapping of protein lysine methylation identifies new functions for HP1β in DNA damage repair. Molecular cell 50(5): 723–735.

Liu, Hongcheng, Ponniah, G., Neill, A., Patel, R. & Andrien, B. (2013) Accurate Determination of Protein Methionine Oxidation by Stable Isotope Labeling and LC-MS Analysis. Analytical Chemistry 85(24): 11705–11709.

Mann, M. & Jensen, O.N. (2003) Proteomic analysis of post-translational modifications. Nature Biotechnology 21(3): 255–261.

Martin, S.C., Ekman, P., Forsberg, P.O. & Ersmark, H. (1992) Increased phosphate content of fibrinogen in vivo correlates with alteration in fibrinogen behaviour. Thrombosis Research 68(6): 467–473.

Maverakis, E., Kim, K., Shimoda, M., Gershwin, M.E., Patel, F., Wilken, R., Raychaudhuri, S., Ruhaak, L.R. & Lebrilla, C.B. (2015) Glycans in the immune system and The Altered Glycan Theory of Autoimmunity: A critical review. Journal of Autoimmunity 57: 1–13.

Mazzon, M., Jones, M., Davidson, A., Chain, B. & Jacobs, M. (2009) Dengue virus NS5 inhibits interferon-alpha signaling by blocking signal transducer and activator of transcription 2 phosphorylation. The Journal of Infectious Diseases 200(8): 1261–1270.

Mi, H., Ebert, D., Muruganujan, A., Mills, C., Albou, L.-P., Mushayamaha, T. & Thomas, P.D. (2021) PANTHER version 16: a revised family classification, tree-based classification tool, enhancer regions and extensive API. Nucleic Acids Research 49(D1): D394–D403.

Mi, H., Muruganujan, A., Huang, X., Ebert, D., Mills, C., Guo, X. & Thomas, P.D. (2019) Protocol Update for large-scale genome and gene function analysis with the PANTHER classification system (v.14.0). Nature Protocols 14(3): 703–721.

Miao, M., Yu, F., Wang, D., Tong, Y., Yang, L., Xu, J., Qiu, Y., Zhou, X. & Zhao, X. (2019) Proteomics Profiling of Host Cell Response via Protein Expression and Phosphorylation upon Dengue Virus Infection. Virologica Sinica 34(5): 549–562.

Minguez, P., Parca, L., Diella, F., Mende, D.R., Kumar, R., Helmer-Citterich, M., Gavin, A.-C., van Noort, V. & Bork, P. (2012) Deciphering a global network of functionally associated post-translational modifications. Molecular Systems Biology 8(1): 599.

Mnatsakanyan, R., Shema, G., Basik, M., Batist, G., Borchers, C.H., Sickmann, A. & Zahedi, R.P. (2018) Detecting post-translational modification signatures as potential biomarkers in clinical mass spectrometry. Expert Review of Proteomics 15(6): 515–535.

Morand, K., Talbo, G. & Mann, M. (1993) Oxidation of peptides during electrospray ionization. Rapid communications in mass spectrometry: RCM 7(8): 738–743.

Nair, R.R., Khanna, A. & Singh, K. (2013) Role of inflammatory proteins S100A8 and S100A9 in pathophysiology of recurrent early pregnancy loss. Placenta 34(9): 824–827.

Ngono, A.E. & Shresta, S. (2019) Cross-Reactive T Cell Immunity to Dengue and Zika Viruses: New Insights Into Vaccine Development. Frontiers in Immunology 10. Available from: https://www.ncbi.nlm.nih.gov/pmc/articles/PMC6579874/ [Accessed 11 June 2020].

Oehmcke-Hecht, S. & Köhler, J. (2018) Interaction of the Human Contact System with Pathogens—An Update. Frontiers in Immunology 9. Available from: https://www.ncbi.nlm.nih.gov/pmc/articles/PMC5834483/ [Accessed 20 February 2021].

Oriá, R.B., Patrick, P.D., Zhang, H., Lorntz, B., de Castro Costa, C.M., Brito, G.A.C., Barrett, L.J., Lima, A.A.M. & Guerrant, R.L. (2005) APOE4 Protects the Cognitive Development in Children with Heavy Diarrhea Burdens in Northeast Brazil. Pediatric Research 57(2): 310–316.

Osuna-Ramos, J.F., Reyes-Ruiz, J.M. & del Ángel, R.M. (2018) The Role of Host Cholesterol During Flavivirus Infection. Frontiers in Cellular and Infection Microbiology 8. Available from: https://www.ncbi.nlm.nih.gov/pmc/articles/PMC6224431/ [Accessed 3 February 2021].

Parekh, R.B., Dwek, R.A., Sutton, B.J., Fernandes, D.L., Leung, A., Stanworth, D., Rademacher, T.W., Mizuochi, T., Taniguchi, T., Matsuta, K., Takeuchi, F., Nagano, Y., Miyamoto, T. & Kobata, A. (1985) Association of rheumatoid arthritis and primary osteoarthritis with changes in the glycosylation pattern of total serum IgG. Nature 316(6027): 452–457.

Pascovici, D., Wu, J.X., McKay, M.J., Joseph, C., Noor, Z., Kamath, K., Wu, Y., Ranganathan, S., Gupta, V. & Mirzaei, M. (2018) Clinically Relevant Post-Translational Modification Analyses—Maturing Workflows and Bioinformatics Tools. International Journal of Molecular Sciences 20(1). Available from: https://www.ncbi.nlm.nih.gov/pmc/articles/PMC6337699/ [Accessed 12 September 2020].

Pederson, E.N. & Interlandi, G. (2019) Oxidation-induced destabilization of the fibrinogen αC-domain dimer investigated by molecular dynamics simulations. Proteins 87(10): 826–836.

Pierson, T.C. & Diamond, M.S. (2020) The continued threat of emerging flaviviruses. Nature Microbiology 5(6): 796–812.

Pierson, T.C., Xu, Q., Nelson, S., Oliphant, T., Nybakken, G.E., Fremont, D.H. & Diamond, M.S. (2007) The stoichiometry of antibody-mediated neutralization and enhancement of West Nile virus infection. Cell Host & Microbe 1(2): 135–145.

Potgieter, H.C., Ubbink, J.B., Bissbort, S., Bester, M.J., Spies, J.H. & Vermaak, W.J. (1997) Spontaneous oxidation of methionine: effect on the quantification of plasma methionine levels. Analytical Biochemistry 248(1): 86–93.

Ray, S., Srivastava, R., Tripathi, K., Vaibhav, V., Patankar, S. & Srivastava, S. (2012) Serum proteome changes in dengue virus-infected patients from a dengue-endemic area of India: towards new molecular targets? Omics : a journal of integrative biology.

Ree, R., Varland, S. & Arnesen, T. (2018) Spotlight on protein N-terminal acetylation. Experimental & Molecular Medicine 50(7): 1–13.

Reiter, L., Rinner, O., Picotti, P., Hüttenhain, R., Beck, M., Brusniak, M.-Y., Hengartner, M.O. & Aebersold, R. (2011) mProphet: automated data processing and statistical validation for large-scale SRM experiments. Nature Methods 8(5): 430–435.

Rosenberger, G., Koh, C.C., Guo, T., Röst, H.L., Kouvonen, P., Collins, B.C., Heusel, M., Liu, Y., Caron, E., Vichalkovski, A., Faini, M., Schubert, O.T., Faridi, P., Ebhardt, H.A., Matondo, M., Lam, H., Bader, S.L., Campbell, D.S., Deutsch, E.W., Moritz, R.L., Tate, S. & Aebersold, R. (2014) A repository of assays to quantify 10,000 human proteins by SWATH-MS. Scientific Data 1: 140031.

Sajic, T., Liu, Y. & Aebersold, R. (2015) Using data-independent, high-resolution mass spectrometry in protein biomarker research: perspectives and clinical applications. Proteomics. Clinical Applications 9(3–4): 307–321.

Schaller, J., Gerber, S., Kaempfer, U., Lejon, S. & Trachsel, C. (2008) Posttranslational Modifications. In: Human Blood Plasma Proteins. John Wiley & Sons, Ltd, 75–87. Available from: https://onlinelibrary.wiley.com/doi/abs/10.1002/9780470724378.ch6 [Accessed 11 June 2020].

Schenten, V., Plançon, S., Jung, N., Hann, J., Bueb, J.-L., Bréchard, S., Tschirhart, E.J. & Tolle, F. (2018) Secretion of the Phosphorylated Form of S100A9 from Neutrophils Is Essential for the Proinflammatory Functions of Extracellular S100A8/A9. Frontiers in Immunology 9: 447.

Seo, J. & Lee, K.-J. (2004) Post-translational modifications and their biological functions: proteomic analysis and systematic approaches. Journal of Biochemistry and Molecular Biology 37(1): 35–44.

Shaklai, N., Garlick, R.L. & Bunn, H.F. (1984) Nonenzymatic glycosylation of human serum albumin alters its conformation and function. Journal of Biological Chemistry 259(6): 3812–3817.

Shao, B. (2012) Site-specific oxidation of apolipoprotein A-I impairs cholesterol export by ABCA1, a key cardioprotective function of HDL. Biochimica Et Biophysica Acta 1821(3): 490–501.

Shao, B., Cavigiolio, G., Brot, N., Oda, M.N. & Heinecke, J.W. (2008) Methionine oxidation impairs reverse cholesterol transport by apolipoprotein A-I. Proceedings of the National Academy of Sciences of the United States of America 105(34): 12224–12229.

Sharma, S. (2021) India’s latest Zika outbreak sees surge of nearly 100 cases. Reuters. Available from: https://www.reuters.com/business/healthcare-pharmaceuticals/indias-latest-zika-outbreak-sees-surge-nearly-100-cases-2021-11-08/ [Accessed 9 November 2021].

Sharp, T.M., Fischer, M., Muñoz-Jordán, J.L., Paz-Bailey, G., Staples, J.E., Gregory, C.J. & Waterman, S.H. (2019) Dengue and Zika Virus Diagnostic Testing for Patients with a Clinically Compatible Illness and Risk for Infection with Both Viruses. MMWR. Recommendations and reports: Morbidity and mortality weekly report. Recommendations and reports 68(1): 1–10.

Shevchenko, A., Wilm, M., Vorm, O. & Mann, M. (1996) Mass Spectrometric Sequencing of Proteins from Silver-Stained Polyacrylamide Gels. Available from: https://pubs.acs.org/doi/abs/10.1021/ac950914h [Accessed 13 March 2021].

Sidoli, S., a, G.B., r, K.K., Katarzyna, K. & Johayra, S. (2015) Low Resolution Data-Independent Acquisition in an LTQ-Orbitrap Allows for Simplified and Fully Untargeted Analysis of Histone Modifications. Analytical chemistry. Available from: https://agris.fao.org/agris-search/search.do?recordID=US201700032725 [Accessed 29 April 2021].

Spiller, S., Li, Y., Blüher, M., Welch, L. & Hoffmann, R. (2017) Glycated lysine-141 in haptoglobin improves the diagnostic accuracy for type 2 diabetes mellitus in combination with glycated hemoglobin HbA1c and fasting plasma glucose. Clinical Proteomics 14: 10.

Subissi, L., Dub, T., Besnard, M., Mariteragi-Helle, T., Nhan, T., Lutringer-Magnin, D., Barboza, P., Gurry, C., Brindel, P., Nilles, E.J., Baud, D., Merianos, A., Musso, D., Glynn, J.R., Dupuis, G., Cao-Lormeau, V.-M., Giard, M. & Mallet, H.-P. (2018) Zika Virus Infection during Pregnancy and Effects on Early Childhood Development, French Polynesia, 2013–2016. Emerging Infectious Diseases 24(10): 1850–1858.

Sun, S., Hu, Y., Jia, L., Eshghi, S.T., Liu, Y., Shah, P. & Zhang, H. (2018) Site-Specific Profiling of Serum Glycoproteins Using N-Linked Glycan and Glycosite Analysis Revealing Atypical N-Glycosylation Sites on Albumin and α-1B-Glycoprotein. Analytical Chemistry 90(10): 6292–6299.

Taggart, C., Cervantes-Laurean, D., Kim, G., McElvaney, N.G., Wehr, N., Moss, J. & Levine, R.L. (2000) Oxidation of either methionine 351 or methionine 358 in alpha 1-antitrypsin causes loss of anti-neutrophil elastase activity. The Journal of Biological Chemistry 275(35): 27258–27265.

Thulin, N.K., Brewer, R.C., Sherwood, R., Bournazos, S., Edwards, K.G., Ramadoss, N.S., Taubenberger, J.K., Memoli, M., Gentles, A.J., Jagannathan, P., Zhang, S., Libraty, D.H. & Wang, T.T. (2020) Maternal Anti-Dengue IgG Fucosylation Predicts Susceptibility to Dengue Disease in Infants. Cell Reports 31(6): 107642.

Varland, S., Osberg, C. & Arnesen, T. (2015) N-terminal modifications of cellular proteins: The enzymes involved, their substrate specificities and biological effects. PROTEOMICS 15(14): 2385–2401.

de Vries, J.J., Snoek, C.J.M., Rijken, D.C. & de Maat, M.P.M. (2020) Effects of Post-Translational Modifications of Fibrinogen on Clot Formation, Clot Structure, and Fibrinolysis: A Systematic Review. Arteriosclerosis, Thrombosis, and Vascular Biology 40(3): 554–569.

Wang, B., Zhu, Y., Yu, C., Zhang, C., Tang, Q., Huang, H. & Zhao, Z. (2021) Hepatitis C virus induces oxidation and degradation of apolipoprotein B to enhance lipid accumulation and promote viral production. PLOS Pathogens 17(9): e1009889.

Wei, B., Berning, K., Quan, C. & Zhang, Y.T. (2017) Glycation of antibodies: Modification, methods and potential effects on biological functions. mAbs 9(4): 586–594.

Weigandt, K.M., White, N., Chung, D., Ellingson, E., Wang, Y., Fu, X. & Pozzo, D.C. (2012) Fibrin clot structure and mechanics associated with specific oxidation of methionine residues in fibrinogen. Biophysical Journal 103(11): 2399–2407.

Wen, J., Tang, W.W., Sheets, N., Ellison, J., Sette, A., Kim, K. & Shresta, S. (2017) Identification of Zika virus epitopes reveals immunodominant and protective roles for dengue virus cross-reactive CD8 + T cells. Nature Microbiology 2(6): 1–11.

White, N.J., Wang, Y., Fu, X., Cardenas, J.C., Martin, E.J., Brophy, D.F., Wade, C.E., Wang, X., St John, A.E., Lim, E.B., Stern, S.A., Ward, K.R., López, J.A. & Chung, D. (2016) Post-translational oxidative modification of fibrinogen is associated with coagulopathy after traumatic injury. Free Radical Biology & Medicine 96: 181–189.

Wong, Y.Q., Binger, K.J., Howlett, G.J. & Griffin, M.D.W. (2010) Methionine oxidation induces amyloid fibril formation by full-length apolipoprotein A-I. Proceedings of the National Academy of Sciences of the United States of America 107(5): 1977–1982.

Wu, S.M., Patel, D.D. & Pizzo, S.V. (1998) Oxidized alpha2-macroglobulin (alpha2M) differentially regulates receptor binding by cytokines/growth factors: implications for tissue injury and repair mechanisms in inflammation. Journal of Immunology (Baltimore, Md.: 1950) 161(8): 4356–4365.

Xu, H., Wang, Y., Lin, S., Deng, W., Peng, D., Cui, Q. & Xue, Y. (2018) PTMD: A Database of Human Disease-associated Post-translational Modifications. Genomics, Proteomics & Bioinformatics 16(4): 244–251.

Yalak, G., Shiu, J.-Y., Schoen, I., Mitsi, M. & Vogel, V. (2019) Phosphorylated fibronectin enhances cell attachment and upregulates mechanical cell functions. PLOS ONE 14(7): e0218893.

Yeom, J., Ju, S., Choi, Y., Paek, E. & Lee, C. (2017) Comprehensive analysis of human protein N-termini enables assessment of various protein forms. Scientific Reports 7(1): 6599.

CDC. 2020. “Dengue Virus.” July 16, 2020. https://www.cdc.gov/dengue/index.html.

